# Langerhans Cells Regulate Tongue Intraepithelial Innervation in a Microbiota- and Age-Dependent Manner

**DOI:** 10.1101/2025.04.03.647024

**Authors:** Yasmin Netanely, Yasmin Saba, Reem Naamneh, Shahd Yacoub, Yasmin Jaber, Alon Raviv, Luba Eli-Berchoer, Hagit Shapiro, Eran Elinav, Asaf Wilensky, Björn E. Clausen, Hovav Avi-Hai

**Author notes:** Co-first authors (Yasmin Netanely, Yasmin Saba).

## Abstract

Oral Langerhans cells (LCs) are well recognized for their immunological roles, but their involvement in other physiological processes remains poorly understood. This study identifies a novel function of oral LCs in regulating tongue epithelial innervation. Postnatal LC development coincides with the establishment of local innervation, and LC depletion impairs innervation and alters nociceptive responses, underscoring their neuroimmune function. This function is driven by LC-derived IL-1β, which stimulates basal epithelial cells to produce nerve growth factor (NGF), thereby promoting sensory nerve growth. Transcriptomic analyses revealed neuronal-related pathways enriched in LCs. Aging reduces LC frequency, NGF expression, and epithelial innervation, linking neuroimmune regulation to epithelial aging. While LC frequencies in the tongue remain unaffected in germ-free mice, the microbiota is essential for optimal LC function and NGF production. These findings expand our understanding of oral LCs, revealing their pivotal role in epithelial innervation beyond immune surveillance.

## Introduction

Langerhans cells (LCs) are specialized antigen-presenting cells (APCs), residing in the stratified squamous epithelia such as the skin and oral mucosa (Brand et al., 2023). Skin LCs primarily arise from embryonic precursors, exhibiting traits of both dendritic cells (DCs) and tissue-resident macrophages with the capacity for self-renewal (Doebel et al., 2017). In contrast, oral LCs are derived and continuously replenished from adult bone marrow (BM) precursors, similar to classical DCs (cDCs) (Capucha et al., 2018; Capucha et al., 2015). The development of oral LCs, particularly those near the dental biofilm, is influenced by the microbiota, whereas skin LCs develop independently of microbial influence (Capucha et al., 2018). Despite these ontogenetic differences, both skin and oral LCs display a shared "LC signature" characterized by the expression of langerin and EpCAM, the presence of Birbeck granules, and their role as immune sentinels in stratified epithelia, capturing and presenting antigens to initiate immune responses (Clausen and Stoitzner, 2015; Hovav, 2018).

In addition to their immune functions, skin LCs exhibit neural-related roles. Skin LCs form close associations with intraepithelial nerve fibers (Doss and Smith, 2012; Gaudillere et al., 1996), and their density in the epidermis is regulated by cutaneous innervation (Hsieh et al., 1996; Siau et al., 2006; Stankovic et al., 1999). Conversely, depletion of LCs leads to reduced epidermal innervation and results in mechanical hypersensitivity in the footpad (Doss and Smith, 2014). Neuropeptides also play a significant role in modulating LC function in the skin. Early studies identified the regulation of LC function by nerves containing calcitonin gene-related peptide (CGRP), which inhibits the induction of delayed-type and contact hypersensitivity responses (Asahina et al., 1995; Hosoi et al., 1993). Further studies revealed that pituitary adenylate cyclase-activating polypeptide (PACAP) and vasoactive intestinal peptide (VIP) are endogenous mediators that regulate skin immunity by modulating LC activity through the inhibition of NF-κB activation (Hosoi et al., 1993; Kodali et al., 2004; Kodali et al., 2003). In addition, CGRP biases LCs toward promoting Th2-type immunity (Ding et al., 2008), while PACAP and VIP guide LC-mediated antigen presentation toward Th17 responses (Ding et al., 2012). These findings emphasize the bidirectional communication between LCs and intraepithelial nerves in the skin, while the nature of such interaction in the oral epithelium remains largely unexplored.

While the neurally related functions of skin LCs align with their role as tissue-resident macrophages (Msheik et al., 2022), oral LCs are generally regarded as lacking these macrophage-like characteristics. This raises the question of whether oral LCs contribute to intraepithelial innervation in the oral mucosa. This is especially relevant in the tongue, where innervation is crucial for sensory perception and tissue homeostasis (Bosma, 1976; Calhoun et al., 1992). The tongue is densely innervated by sensory nerve fibers that are essential for detecting mechanical, chemical, and thermal stimuli, which are vital for functions such as taste, mastication, and nociception (Kumari and Mistretta, 2023). These sensory inputs are not only critical for protective reflexes but also for maintaining mucosal barrier integrity (Mallesh et al., 2022) and regulating mucosal immunity (Fornai et al., 2018). The restricted location of tongue macrophages, particularly CX_3_CR1^+^ macrophages, in the highly innervated lamina propria beneath the epithelium (Lyras et al., 2022), suggests that LCs within the epithelium could also have neurally related functions. Understanding the role of oral LCs in intraepithelial innervation is crucial for elucidating neuroimmune interactions that regulate oral sensory function.

## Results

### The population of tongue LCs coincides with the development of intraepithelial innervation

To investigate the role of LCs in oral intraepithelial innervation, we first characterized the development of LCs in the tongue epithelium after birth. Flow cytometric analysis revealed that LCs began to populate the epithelium before weaning, with their frequencies at 3 weeks of age (weaning) comparable to those observed in adult mice (Fig. 1A). Immunofluorescence staining confirmed the presence of MHCII^+^ and langerin^+^ LCs in the epithelium postnatally, and their absolute numbers increased as the mice grew and matured into adulthood (Fig. 1B). LCs were exclusively located in the epithelium as no Langerin^+^EpCAM^+^ cells were detected in the underlying lamina propria (Fig. S1). The generation of chimeric mice in which CD45.1^+^ BM cells were transplanted into lethally irradiated CD45.2^+^ recipients, indicated that tongue LCs differentiated from adult BM precursors (Fig. 1C). We then characterized the expression of TGF-β1 and BMP7, two factors known to regulate LC differentiation in the oral epithelium (Capucha et al., 2018). As shown in Fig. 1D, both TGF-β1 and BMP7 were upregulated postnatally, with TGF-β1 localized in the epithelium and BMP7 in the lamina propria. Notably, mRNA levels of both genes increased before and during weaning, aligning with LC development, and subsequently returned to baseline levels (Fig. 1E). To visualize peripheral nerve fibers, we stained tongue cross-sections with antibodies against the pan-neuronal markers PGP9.5 and β3-tubulin. While staining with anti-PGP9.5 antibody did not reveal free nerve endings in the tongue epithelium of 1 week-old mice, these became evident by the fourth week, and the epithelium was highly innervated by 8 weeks of age (Fig. 1F). In addition, co-staining with MHCII antibodies revealed MHCII-positive cells in the lamina propria of 1 week-old mice, likely macrophages or cDCs, that colocalized with nerve fibers in a region that was already innervated at this early age (Fig. 1F). In the epithelium, MHCII-positive cells were observed in 4 week-old mice (representing LCs), and their numbers increased by 8 weeks of age, when they were located close to peripheral nerves. Staining with β3-tubulin antibody further confirmed the gradual development of intraepithelial innervation after birth (Fig. 1G). These findings highlight the concurrent development of LCs and innervation in the tongue epithelium after birth.

**Figure 1:**
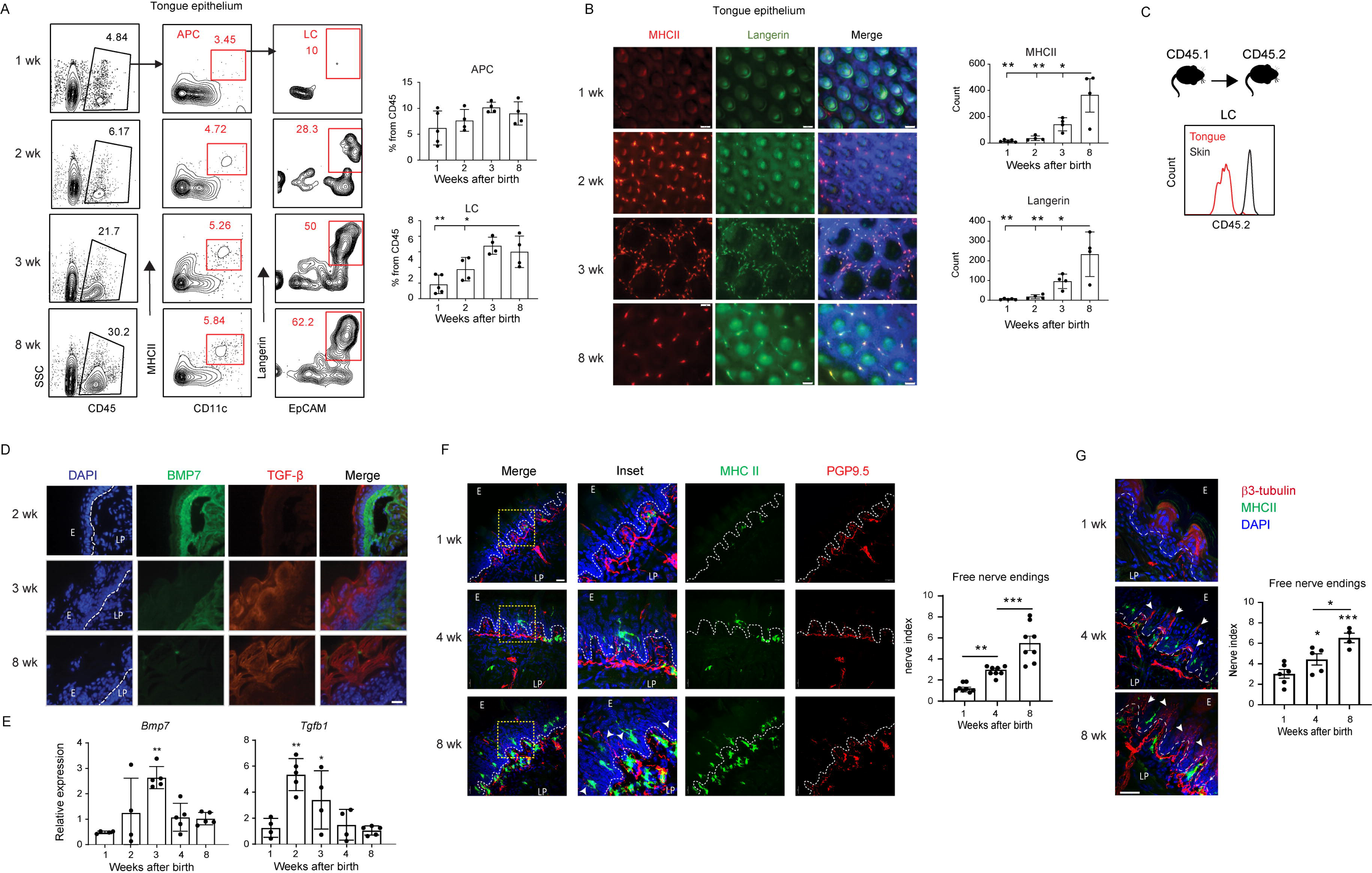
Postnatal development of LCs and innervation in the tongue epithelium **(A)** Representative flow cytometry plots and graphs showing the mean frequencies ± SEM (n=4-5) of APCs and LCs in the tongue epithelium from birth to adulthood. Data are representative of three independent experiments. **(B)** Whole-mount immunofluorescence images of tongue epithelial sheets at various postnatal weeks, stained with mAbs against MHCII (red), langerin (green), and DAPI (blue) for nuclear visualization. Graphs show the mean numbers of MHCII⁺ or langerin⁺ cells ± SEM (n=4-5). Data are representative of two independent experiments. Scale bar, 50 µm. **(C)** Flow cytometry histogram showing CD45.2 expression on tongue and skin LCs from lethally irradiated CD45.2⁺ B6 mice transplanted with BM purified from CD45.1⁺ B6 mice. Data are representative of two independent experiments. **(D)** Immunofluorescence images of tongue cross-sections at various postnatal weeks, stained for TGF-β1 (red), BMP7 (green), and DAPI (blue). The white line indicates the basal membrane. E, epithelium; LP, lamina propria. Data are representative of two independent experiments. Scale bar, 50 µm. **(E)** Relative expression levels of *Tgfb1* and *Bmp7* in the tongue at various postnatal weeks. Graphs display mean transcript levels ± SEM (n=4-5), quantified by qPCR and normalized to 8-week-old mice. **(F-G)** Immunofluorescence images of tongue cross-sections at various postnatal weeks, stained for PGP9.5 or β3-tubulin (red), MHCII (green), and DAPI (blue). White dotted lines indicate the basal membrane. Graphs display the mean nerve index ± SEM (n=4-8). Data are representative of two independent experiments. Scale bar, 50 µm. *, P < 0.05, **, P < 0.01, and *** P < 0.00.1.

### Conditional depletion of LCs reduces intraepithelial innervation in the tongue and impairs the nociceptive pain response

To investigate whether oral LCs regulate epithelial innervation, we utilized Langerin-DTR mice, which allow for the conditional ablation of langerin-expressing cells upon diphtheria toxin (DT) administration (Bennett et al., 2005). First, we assessed the depletion kinetics of tongue LCs, demonstrating that following a single DT injection, LCs were rapidly depleted but completely replenished within 6 days (Fig. 2A). Notably, LC repopulation occurred significantly faster in the tongue compared to gingival LCs (Fig. 2B), as EpCAM^+^MHCII^+^ cells, representing immediate LC precursors, swiftly migrated into the epithelium (Fig. 2A, C). This was not due to an increased proliferation rate of tongue LCs, since gingival and tongue LCs have similar BrdU incorporation kinetics (Fig. S2). As expected, LCs did not repopulate the skin epidermis within the observed period (Bennett et al., 2005). Given this rapid repopulation rate, DT, and PBS as a negative control, were administered twice weekly to Langerin-DTR mice, starting 2 weeks after birth to encompass the critical period of intraepithelial innervation establishment (Fig. 2D). Analysis of LC-depleted mice at 8 weeks of age showed a marked reduction in PGP9.5 and β3-tubulin staining of intraepithelial innervation within the tongue epithelium, while the innervation of the lamina propria was unaffected (Fig. 2D). Notably, stopping DT injections in adult LC-depleted mice restored intraepithelial innervation within 3 weeks (Fig. 2E). Additionally, administering DT to adult Langerin-DTR mice for 3 weeks also led to reduced intraepithelial innervation (Fig. 2F). Finally, we assessed aversion to capsaicin in drinking water in postnatally DT-treated mice. As depicted in Fig. 2G, LC-depleted mice consumed the water normally, while LC-intact (PBS-treated) mice avoided it (see Videos 1 and 2). These findings demonstrate that LCs regulate intraepithelial innervation in the tongue and modulate nociceptive responses. The restoration of innervation after LC depletion ceases suggests that LCs sustain nerve presence or growth without affecting neuronal viability.

**Figure 2:**
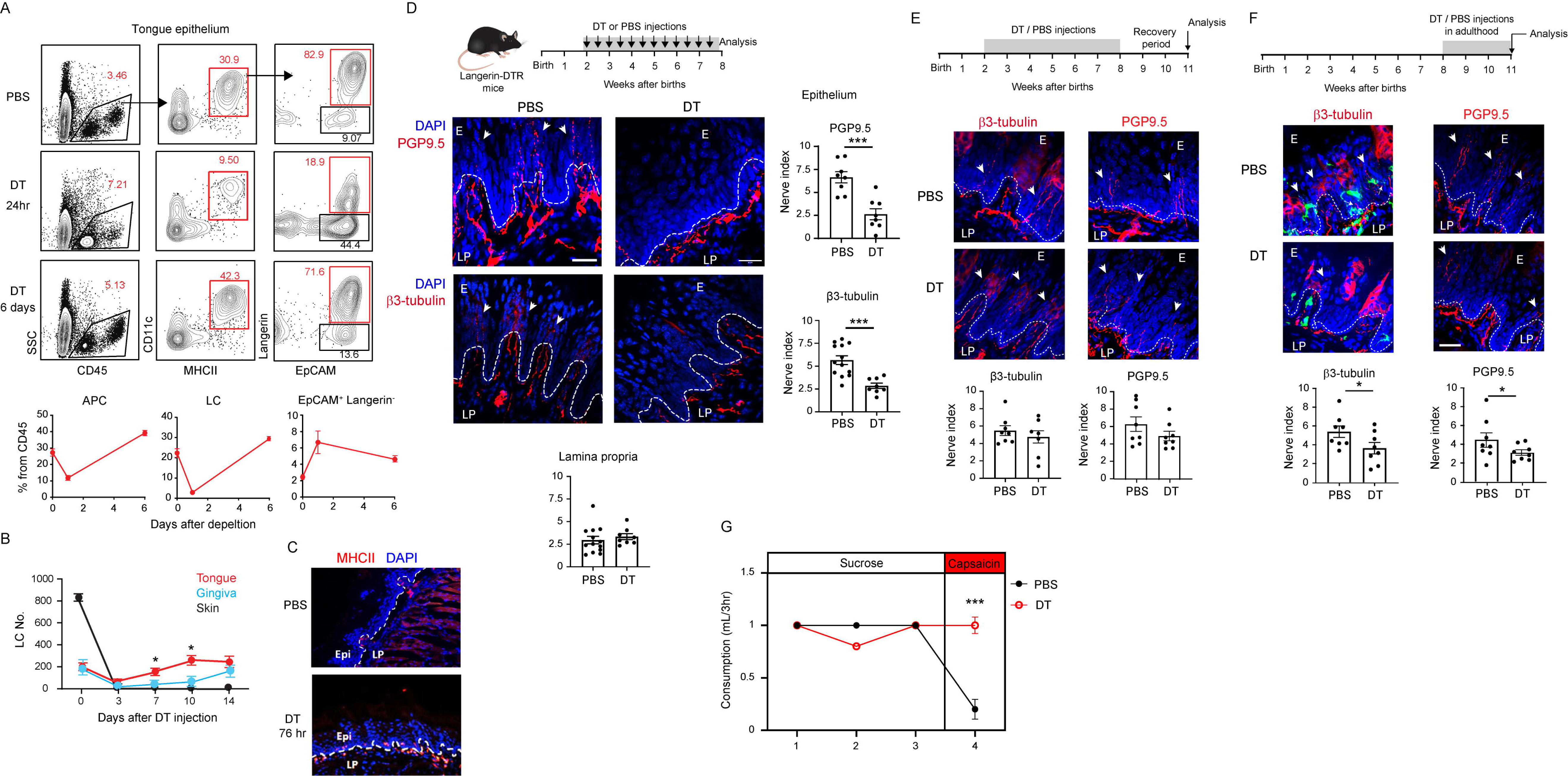
Depletion of tongue LCs reduces intraepithelial innervation and impairs nociceptive pain response **(A)** Representative flow cytometry plots and graphs showing the mean frequencies ± SEM (n=5) of APCs, LCs, and EpCAM⁺ DCs in the tongue epithelium of Langerin-DTR mice at various time points after DT injection. Data are representative of three independent experiments. **(B)** Repopulation kinetics of LCs in the tongue, gingiva, and skin following DT injections, presented as mean frequencies ± SEM (n=5). **(C)** Immunofluorescence images of tongue cross-sections 76 hours post-DT or PBS injection, stained for MHCII (red) and DAPI (blue). Data are representative of two independent experiments. The white dotted line indicates the basal membrane. **(D)** Langerin-DTR mice received DT or PBS as depicted in the figure. Immunofluorescence images of tongue cross-sections from DT- or PBS-treated mice stained for PGP9.5 or β3-tubulin (red) and DAPI (blue). Graphs display the mean nerve index ± SEM (n=8-13). Data are representative of two independent experiments. Scale bar, 50 µm. **(E)** Langerin-DTR mice treated with DT or PBS as in (D) underwent a 3-week recovery period before analysis. Immunofluorescence images of tongue cross-sections stained for PGP9.5 or β3-tubulin (red) and DAPI (blue). Graphs show the mean nerve index ± SEM (n=8). Data are representative of three independent experiments. Scale bar, 50 µm. **(F)** Adult Langerin-DTR mice were administered DT or PBS twice weekly. Tongue cross-sections were stained for PGP9.5, β3-tubulin, and DAPI. Graphs show the mean nerve index ± SEM (n=8). Data are representative of two independent experiments. Scale bar, 50 µm. **(G)** Langerin-DTR mice received DT or PBS twice weekly from 2 weeks of age to adulthood. Mice were then given sucrose-supplemented water for 3 days, followed by capsaicin-supplemented water. Graphs present water consumption ± SEM (n=5). Data are representative of three independent experiments. *, P < 0.05 and *** P < 0.00.1.

### Depletion of LCs reduces NGF expression in the tongue epithelium

Given that nerve growth factor (NGF) plays a key role in supporting sensory neuron survival, differentiation, and maintenance (Barker et al., 2020; Meltzer et al., 2021), we investigated whether LCs regulate NGF expression. Initially, we characterized NGF expression in the tongue epithelium postnatally. As shown in Fig. 3A, both the pro-NGF and mature NGF forms were detected in the epithelium by immunofluorescence staining in tongue cross-sections from 1 week-old mice. NGF expression increased during the weaning period and returned to baseline levels in adulthood. In contrast, qPCR analysis of *Ngf* gene expression showed a gradual increase from birth to adulthood (Fig. 3B), suggesting that distinct post-transcriptional and post-translational mechanisms regulate NGF protein levels. Next, DT was administered to Langerin-DTR mice after birth, as described in Fig. 2A, and tongue cross-sections were prepared from adult mice. NGF expression, including both pro-NGF and mature NGF, was significantly reduced in the epithelium following long-term LC depletion (Fig. 3C). Western blot analysis of epithelial sheets isolated from the tongue (Fig. 3D), and qPCR analysis (Fig. 3E), further confirmed the reduction of NGF expression in DT-treated mice. To better understand the type of epithelial cells producing NGF, we took advantage of our recently published single-cell RNA sequencing (scRNA-seq) dataset of oral epithelial cells (Jaber et al., 2024). As shown in Fig. 3F, NGF mRNA was predominantly detected in basal epithelial cells. To directly confirm NGF expression by epithelial cells, we used flow cytometry to sort the tongue epithelium into epithelial cells (CD45^neg^) and leukocytes (CD45⁺), followed by RNA purification from each population. As shown in Fig. 3G, qPCR analysis revealed high levels of *Ngf* expression in the epithelial cell fraction, but not in the leukocyte fraction. These results indicate that NGF is expressed by tongue epithelial cells and suggest that LCs regulate this expression.

**Figure 3:**
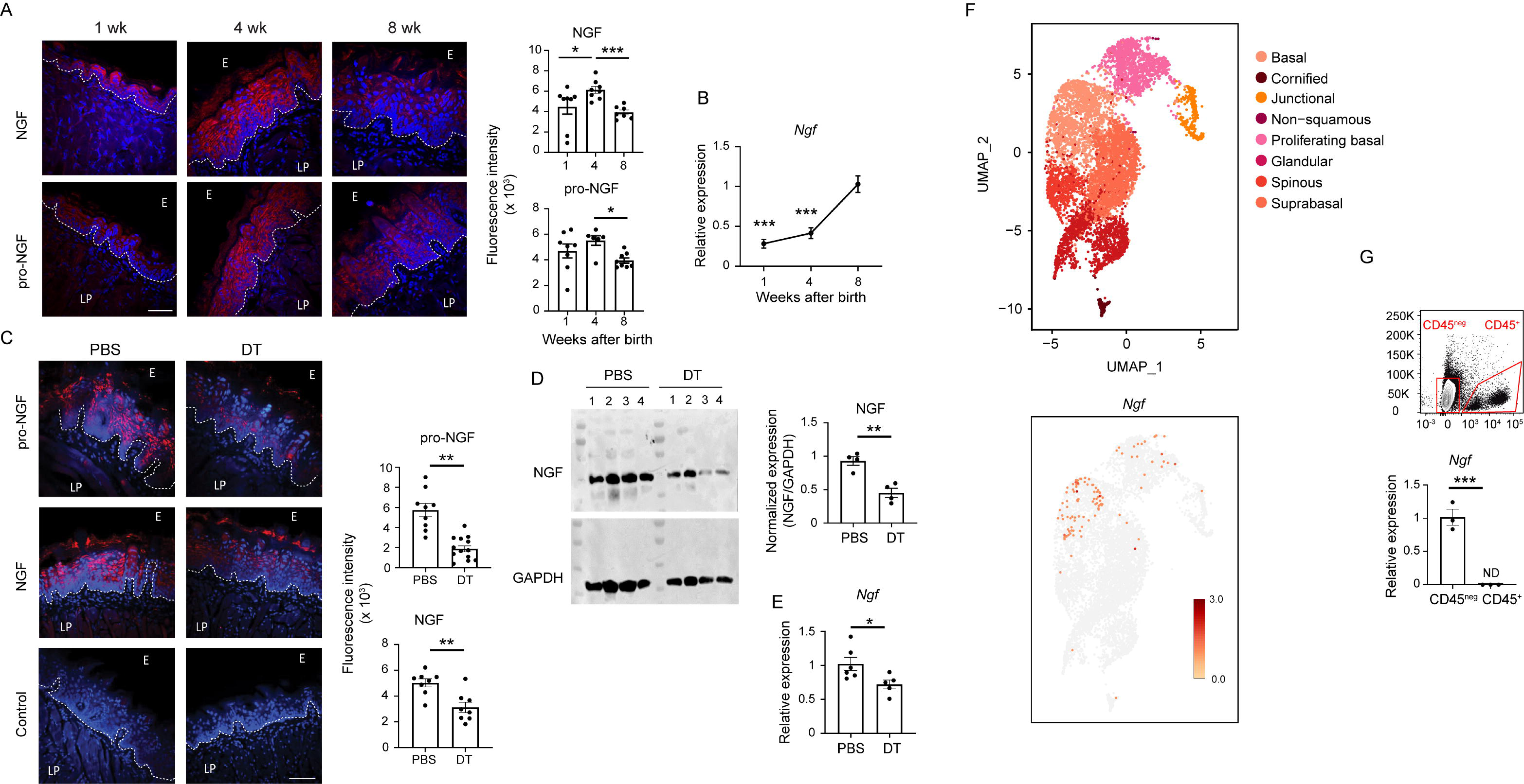
LCs regulate NGF expression in the tongue epithelium. **(A)** Immunofluorescence images of tongue cross-sections at various postnatal weeks, stained for pro-NGF or NGF (red) and DAPI (blue). Graphs display the mean fluorescence intensity ± SEM (n=6-8). Data are representative of two independent experiments. White dotted line indicates the basal membrane. Scale bar, 50 µm. **(B)** Relative expression of the *Ngf* gene in the tongue epithelium at various postnatal weeks. Graphs show mean transcript levels ± SEM (n=5-6), quantified by qPCR and normalized to 8 week-old mice. Data are representative of two independent experiments. **(C–E)** Langerin-DTR mice were administered DT or PBS twice weekly from 2 weeks of age to adulthood. **(C)** Tongue cross-sections stained for pro-NGF or NGF (red) and DAPI (blue). Graphs display mean fluorescence intensity ± SEM (n=8-13). Data are representative of two independent experiments. Scale bar, 50 µm. **(D)** Western blot analysis of NGF in the tongue epithelium of DT- and PBS-treated mice, with GAPDH used as a loading control. Graphs show NGF levels normalized to GAPDH ± SEM (n=4). Data are representative of two independent experiments. **(E)** Relative expression of the *Ngf* gene in the tongue epithelium of DT- and PBS-treated mice. Graphs display mean transcript levels ± SEM (n=5-6). Data are representative of two independent experiments. **(F)** UMAP visualization of major cell types in the oral mucosa expressing epithelial-related genes. Additional UMAP plots show *Ngf* expression in epithelial cells of the oral mucosa. **(G)** Relative expression of the *Ngf* gene in flow cytometry-sorted CD45⁻ and CD45⁺ cells from the tongue epithelium. Graphs display mean transcript levels ± SEM (n=3). Data are representative of two independent experiments. *, P < 0.05, **, P < 0.01, and *** P < 0.00.1.

### IL-1β secreted by LCs regulates NGF expression

To gain insight into the mechanism by which LCs modulate NGF and subsequently intraepithelial innervation in an unbiased manner, we profiled the global gene expression of tongue epithelium by RNA sequencing (RNAseq). The hierarchical clustering and principal component analysis (PCA) indicated a significant difference between the epithelial cells purified from LC-depleted and nondepleted mice (Fig. 4A-B). Gene set enrichment analysis (GSEA) revealed changes in various pathways (Fig. 4C). The only pathway upregulated in the DT group was the IFN-α response pathway, which is associated with peripheral neuropathy (Ekstein et al., 2005). Regarding the downregulated pathways, the top reduced pathways were IL-1 signaling, known to be involved in nerve growth (Peng et al., 2020) and nerve regeneration after injury (Wu et al., 2019). Other downregulated pathways included C-type lectin receptors, MHCII antigen presentation, FcεR signaling, and mTOR signaling, all of which are related to the absence of LCs from the epithelium. The prominent impact of LC absence on IL-1 signaling together with its ability to induce NGF secretion by various cell types (Frossard et al., 2005; Jauneau et al., 2006) encouraged us to further examine the role of IL-1β. First, quantification of IL-1β expression in the tongue epithelium revealed a gradual upregulation in the expression from birth to adulthood (Fig. 4D), resembling the expression of NGF shown earlier (Fig. 3B). Intracellular staining using flow cytometry further revealed that IL-1β is expressed by tongue LCs and also by their immediate epithelial precursors EpCAM^+^langerin^-^ DCs (Fig. 4E). No expression was found by other, i.e. non-antigen-presenting myeloid cells (CD11b^+^CD11c^-^MHCII^-^ cells). Analysis of the expression of IL-1β receptor (IL-1R) by flow cytometry demonstrated that tongue epithelial cells as well as LCs were capable of expressing IL-1R1 (Fig. 4F). Next, to assess directly the role of IL-1β, anti-IL-1β or isotype control antibodies were administered intraperitoneally into mice twice a week from 2 to 8 weeks of age. As depicted in Fig. 4G, staining of tongue cross-sections with PGP9.5 or β3-tubulin revealed a significantly decreased epithelial innervation in anti-IL-1β treated mice compared to treatment with the isotype control. The innervation was also reduced in the lamina propria. Furthermore, administration of anti-IL-1β also decreased the expression of pro-NGF and NGF in the tongue epithelium (Fig. 4H). To demonstrate the role of IL-1β *in vitro*, primary tongue epithelial cells were prepared and cultured with recombinant IL-1β for 76 hr. qPCR analysis demonstrated that the addition of IL-1β upregulated *Ngf* gene expression, whereas *Il1r* was not significantly altered (Fig. 4I). These findings underscore the critical role of IL-1β secreted by LCs in regulating NGF expression.

**Figure 4:**
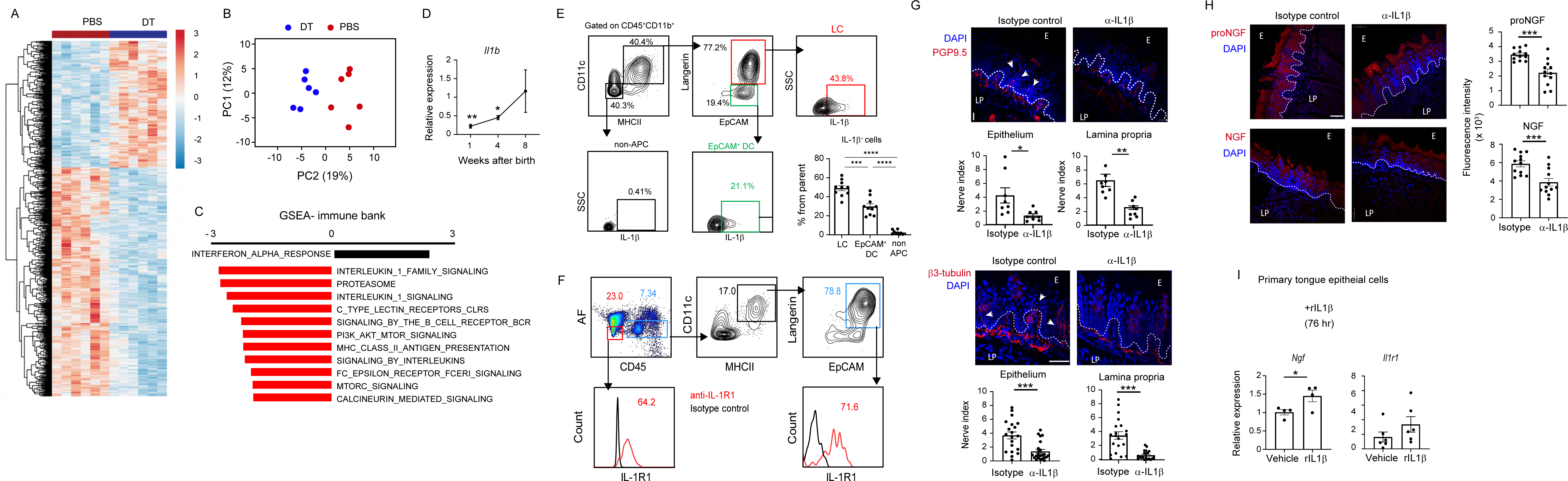
Tongue LCs regulate NGF expression via IL-1β production (A–C) Langerin-DTR mice were administered DT or PBS twice weekly from 2 weeks of age to adulthood. **(A)** Hierarchical clustering and **(B)** principal component analysis (PCA) of genes differentially expressed in the tongue epithelium of DT- and PBS-treated mice. **(C)** Gene set enrichment analysis (GSEA) identifies significantly upregulated and downregulated pathways in DT-versus PBS-treated mice (*FWER* < 0.05). **(D)** Relative expression of the *Il1b* gene in the tongue epithelium at various postnatal weeks. Graph shows mean transcript levels ± SEM (n=5-6), quantified by qPCR and normalized to 8 week-old mice. Data are representative of two independent experiments. **(E)** Flow cytometry plots and graphs showing the mean frequencies ± SEM (n=10) of IL-1β-producing LCs, EpCAM⁺ DCs, and non-APCs in the tongue epithelium. Data are representative of two independent experiments. **(F)** Flow cytometry plots and histograms showing IL-1R1 expression on LCs and tongue epithelial cells. Data are representative of two independent experiments. **(G, H)** B6 mice were administered anti-IL-1β or isotype control antibodies twice weekly from 2 to 8 weeks of age. **(G)** Immunofluorescence images of tongue cross-sections stained for PGP9.5 or β3-tubulin (red) and DAPI (blue). Graphs show the mean nerve index ± SEM (n=8-20). Scale bar, 20 µm and 50 µm, respectively. **(H)** Tongue cross-sections were stained for pro-NGF or NGF (red) and DAPI (blue). Graphs display mean fluorescence intensity ± SEM (n=12). Data are representative of two independent experiments. Scale bar, 50 µm. **(I)** Relative expression of *Ngf* and *Il1r1* genes in primary cultures of tongue epithelial cells incubated with IL-1β for 76 hours. Graphs present mean transcript levels ± SEM (n=4-6). Data are representative of two independent experiments. *, P < 0.05, **, P < 0.01, and *** P < 0.00.1.

### Transcriptomic analysis reveals a neuronal-related signature in tongue LCs

To further investigate tongue LCs, we compared their phenotype with that of the well-characterized gingival LCs. As shown in Fig. 5A, tongue LCs express higher levels of CD11b, CX_3_CR1, and CD64 compared to gingival LCs, indicating a greater contribution from monocytic precurors in agreement with a previous report (Saba et al., 2022). We then reanalyzed published scRNA-seq data from tongue cells (Lyras et al., 2022), identifying LCs as CD74^+^ APCs (excluding B cells) with high expression of langerin and EpCAM (Fig. 5B). Other identified subsets include CX_3_CR1^+^ macrophages, FOLR2^+^ macrophages, Fibronectin 1 (Fn1)^+^ APCs, cDC1 (*Xcr1*), cDC2 (*Irf4^+^irf8^-^*), and *Ly6C*^+^ DCs. Next, we assessed neuronal-related signaling in tongue LCs compared to CX_3_CR1^+^ macrophages, which are known to associate with innervation in the tongue lamina propria and the skin dermis (Kolter et al., 2019; Lyras et al., 2022). As shown in Fig. 5C, while neuroinflammation signaling was more prominent in CX_3_CR1^+^ macrophages, pathways like synaptogenesis, netrin signaling, neuronal morphogenesis, and neurite growth were more enriched in LCs. Both cell types exhibited elevated axonal guidance signaling. Additionally, CX_3_CR1^+^ macrophages showed higher activity in phagosome formation, IL-10, and Toll-like receptor signaling, whereas LCs demonstrated increased pathways related to IL-1 family and TGF-β signaling. We further analyzed neuronal activity among the various mononuclear phagocytes of the tongue. As demonstrated in Fig. 5D, LCs resemble macrophages rather than DCs in pathways associated with the regulation of neurogenesis; but resemble DC in activity associated with the regulation of neuron apoptotic process, an activity predominantly performed by macrophages (Liu et al., 2019). Next, we examined the expression of *Il1b*, *Nen1* (Netrin-1), and *Ngf* across different mononuclear cell populations in the tongue (Fig. 5E). *Il1b* was predominantly expressed by CX_3_CR1^+^ macrophages, Fn1^+^ APCs, cDC2s, and a subset of LCs. In contrast, *Nen1* was primarily expressed by LCs and cDC subsets, while *Ngf* was not expressed by any cell population. Notably, within the LC population, expression of *Il1b* was mainly located in cells expressing *Cxr3r1*, representing monocytic LCs (moLCs) (Fig. 5F). Flow cytometric analysis further substantiated this observation as moLCs (CX_3_CR1^+^ LCs) express higher levels of IL-1β than non-moLCs (Fig. 5F). These findings suggest that tongue LCs play a significant role in neuronal signaling and may interact closely with the nervous system.

**Figure 5:**
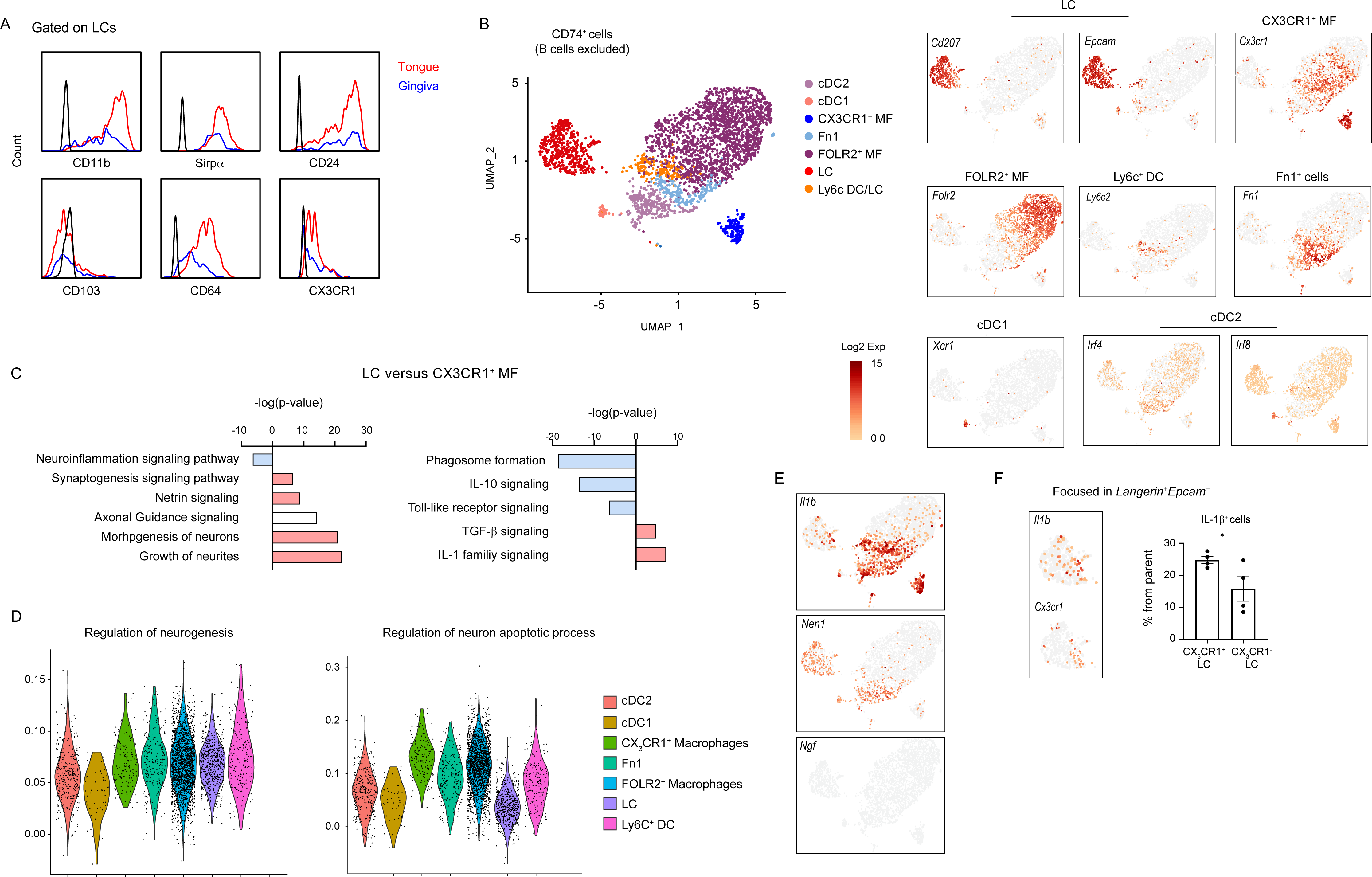
Transcriptomic analysis reveals neuronal-related signature in tongue LCs. **(A)** Representative flow cytometry histograms showing the expression of various phenotypic markers on LCs from the tongue and gingiva. **(B)** UMAP visualization of major mononuclear phagocytes (CD74⁺ cells excluding B cells) in the tongue, based on scRNA-seq data. UMAP plots also display the expression of selected key genes associated with different mononuclear phagocyte subsets. **(C)** Ingenuity Pathway Analysis (IPA) comparing LCs and CX3CR1⁺ macrophages. - log(p-value): red (positive *-log(p-value)*), white (zero *-log(p-value)*), and blue (negative *-log(p-value)*), indicating pathway activation or inhibition. (**D**) Violin plots illustrate the involvement of tongue mononuclear phagocytes in the noted neuronal-related pathways. **(E)** UMAP plots showing the expression of *Il1b*, *Nenl*, and *Ngf* genes in tongue mononuclear phagocytes. (**F**) A graph showing the mean frequencies ± SEM (n=4) of IL-1β-producing moLCs and non-moLC in the tongue epithelium. Data are representative of two independent experiments.

### Aging is associated with reduced LC frequencies and decreased innervation in the tongue epithelium

To examine the impact of aging on LC homeostasis and innervation in the tongue epithelium, we conducted a flow cytometric analysis of 18-month-old and 2-month-old mice. A significant reduction in the frequencies of APCs and LCs was observed in older mice (Fig. 6A). Immunofluorescence staining of tongue epithelial sheets with anti-MHCII and langerin antibodies confirmed this decrease, showing fewer MHCII⁺ langerin⁺ cells in aged mice compared to young controls (Fig. 6B). Additionally, *Tgfb1* expression levels were reduced in older mice, while Bmp7 expression appeared to be upregulated, although not statistically significant (Fig. 6C). We next evaluated local innervation by staining with PGP9.5 and β3-tubulin, observing a reduced nerve index in the tongue epithelium of older mice (Fig. 6D). Furthermore, anti-NGF staining of tongue cross-sections revealed decreased levels of pro-NGF and NGF in aged mice (Fig. 6E). These findings were corroborated by qPCR analysis, which showed diminished *Ngf* expression, along with reduced *Il1b* and *Il1r1* expression, in the tongue epithelium of older mice (Fig. 6F). To investigate the impact of aging on the tongue epithelium, RNAseq analysis was conducted on epithelial sheets isolated from 18-month-old and 2-month-old mice. The hierarchical clustering and PCA indicated a significant difference between the epithelial cells purified from old and young mice (Fig. 6G, H). The GSEA further revealed changes in various pathways (Fig. 6I): Upregulation of the mitotic spindle indicates increased cell division and proliferation in the aged tongue epithelium. The reduction of unfolded protein response, DNA repair, ROS, OXPHOS, and cholesterol homeostasis reflect a compromised cellular environment in aged tongue epithelium, characterized by reduced metabolic and stress-handling capacities. These findings collectively highlight how aging leads to dysregulated epithelial function, characterized by reduced LC frequencies and impaired innervation.

**Figure 6:**
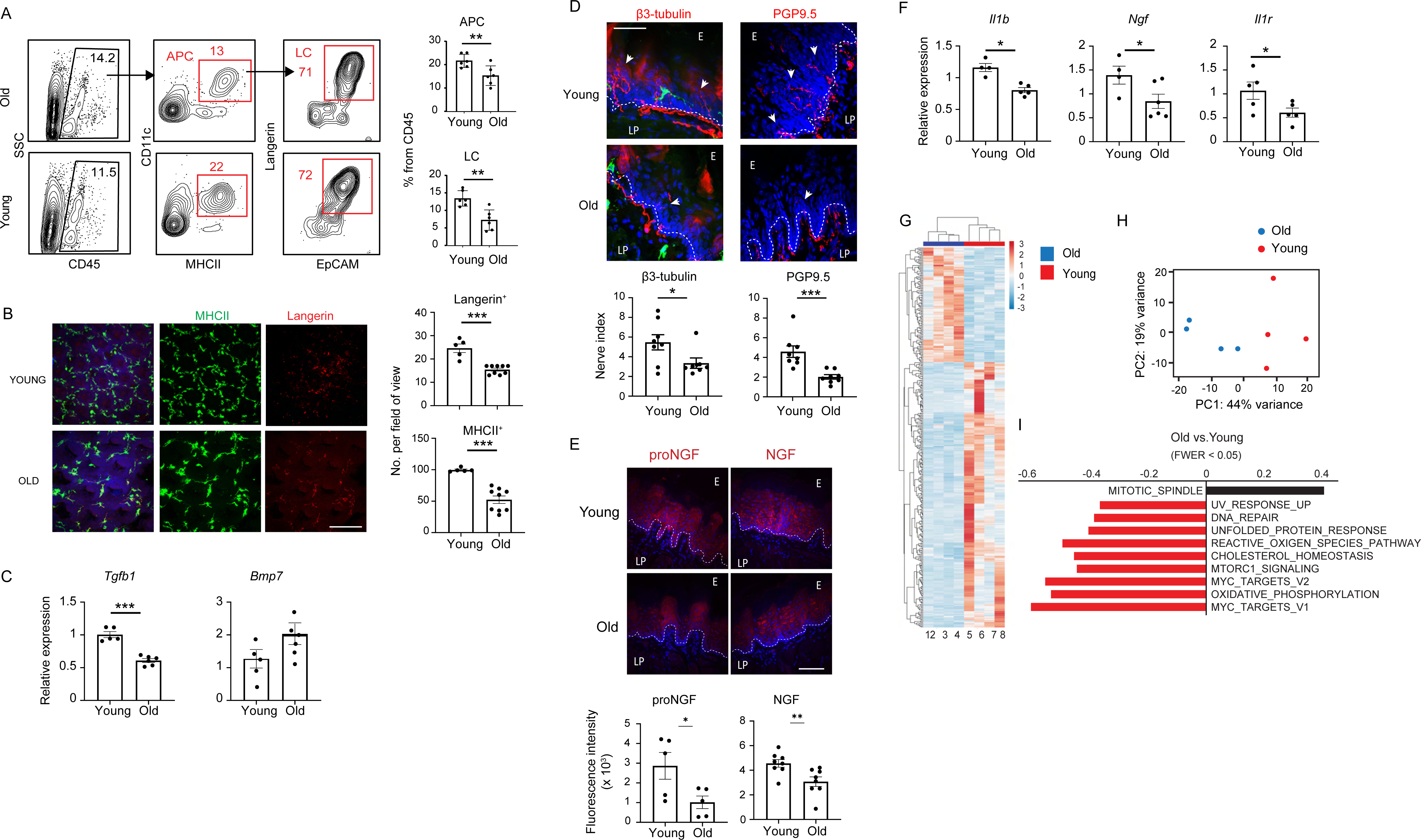
Reduced LCs and decreased innervation in the tongue epithelium with age. **(A)** Representative flow cytometry plots and graphs showing the mean frequencies ± SEM (n=6) of APCs and LCs in the tongue epithelium of 2-month-old (young) and 18-month-old (old) mice. Data are representative of two independent experiments. **(B)** Wholemount immunofluorescence images of tongue epithelial sheets from young and old mice, stained with antibodies against MHCII (red), langerin (green), and DAPI (blue). Graphs display the mean numbers of MHCII⁺ or langerin⁺ cells ± SEM (n=4-5). Data are representative of two independent experiments. Scale bar, 50 µm. **(C)** Relative expression of *Tgfb1* and *Bmp7* genes in tongues of young and old mice. Graphs show mean transcript levels ± SEM (n=5-6), quantified by qPCR. Data are representative of two independent experiments. **(D)** Immunofluorescence images of tongue cross-sections from young and old mice, stained for PGP9.5 or β3-tubulin (red) and DAPI (blue). The white dotted line indicates the basal membrane. Graphs present the mean nerve index ± SEM (n=8). Data are representative of two independent experiments. Scale bar, 50 µm. **(E)** Immunofluorescence images of tongue cross-sections from young and old mice, stained for pro-NGF or NGF (red) and DAPI (blue). Graphs display mean fluorescence intensity ± SEM (n=5-8). Data are representative of two independent experiments. Scale bar, 50 µm. **(F)** Relative expression of selected genes in the tongues of young and old mice. Graphs show mean transcript levels ± SEM (n=4-6), quantified by qPCR. Data are representative of two independent experiments. **(G)** Hierarchical clustering of differentially expressed genes in the tongue epithelium of young and old mice. **(H)** Principal component analysis (PCA) of tongue epithelial gene expression in young and old mice. **(I)** Gene set enrichment analysis (GSEA) identifies significantly upregulated and downregulated pathways between young and old mice (*FWER* < 0.05). *, P < 0.05, **, P < 0.01, and *** P < 0.00.1.

### The microbiota regulates innervation in the tongue epithelium

To investigate the role of the microbiota in intraepithelial innervation of the tongue, we compared germ-free (GF) to specific pathogen-free (SPF) mice. Immunostaining with PGP9.5 and β3-tubulin antibodies revealed diminished innervation in GF mice (Fig. 7A). Notably, while innervation of the lamina propria also appeared to be reduced in GF mice, this difference was not statistically significant. Additionally, Western blot analysis indicated that NGF, but not pro-NGF, levels were decreased in GF mice as compared to SPF controls, which was also confirmed by qPCR (Fig. 7B, C). We then assessed the frequency of LCs in the tongue epithelium. This exhibited no significant differences between GF and SPF mice (Fig. 7D), suggesting that LC development occurs independently of the microbiota. On the other hand, microbial signals might influence the function of the tongue epithelium and local LCs. Supporting this hypothesis, genes involved in TLR signaling, including *Tlr4*, *Tlr2*, and *Myd88*, were downregulated in the tongue epithelium of GF mice (Fig. 7E). Additionally, the expression of *Tgfb1* mRNA (Fig. 7E) and GAS6 protein, a microbiota-dependent homeostatic regulator of the oral epithelium (Nassar et al., 2017), was reduced in GF mice (Fig. 7F). To explore functional changes in LCs, we examined IL-1β production by LCs, revealing reduced expression in GF as compared to SPF mice (Fig. 7G). To confirm the role of the microbiota in regulating LC function, BM-derived LCs were differentiated using GM-CSF and TGF-β1 and stimulated with LPS to activate TLR signaling. As depicted in Fig. 7H, LPS stimulation significantly increased IL-1β production in BM-derived LCs. In summary, although the microbiota does not influence LC frequencies in the tongue epithelium, it is crucial for optimal IL-1β production by LCs via TLR signaling regulation.

**Figure 7:**
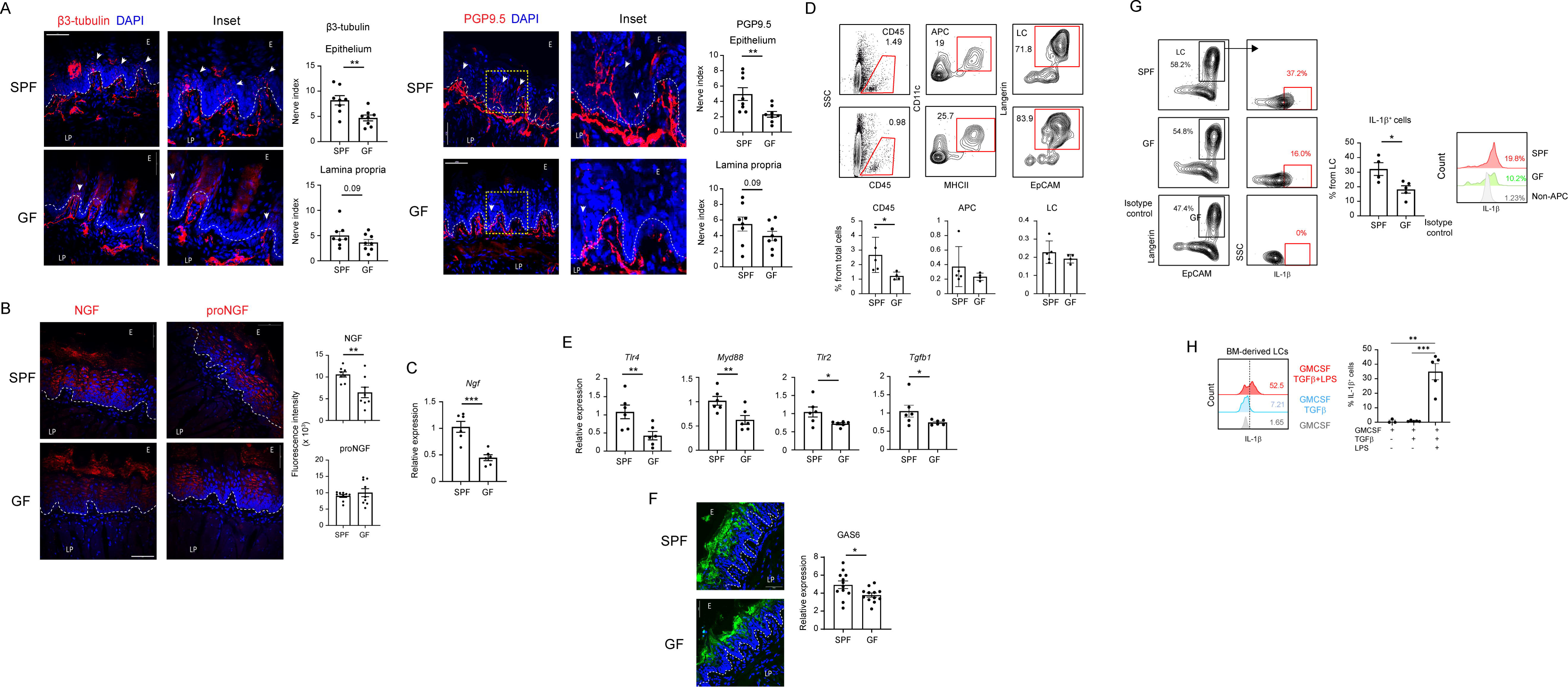
Regulation of tongue intraepithelial innervation by the microbiota. **(A)** Immunofluorescence images of tongue cross-sections from germ-free (GF) and specific pathogen-free (SPF) mice stained for PGP9.5 or β3-tubulin (red) and DAPI (blue). The white dotted line indicates the basal membrane. Graphs display the mean nerve index ± SEM (n=8). Data are representative of two independent experiments. Scale bar, 50 µm. **(B)** Immunofluorescence images of tongue cross-sections from GF and SPF mice stained for pro-NGF or NGF (red) and DAPI (blue). Graphs show mean fluorescence intensity ± SEM (n=8). Data are representative of two independent experiments. Scale bar, 50 µm. **(C)** Relative expression of the *Ngf* gene in the tongue epithelium of GF and SPF mice. Graphs present the mean transcript levels ± SEM (n=6), quantified by qPCR. Data are representative of two independent experiments. **(D)** Representative flow cytometry plots and graphs showing the mean frequencies ± SEM (n=4-5) of CD45⁺ cells, APCs, and LCs in the tongue epithelium of GF and SPF mice. Data are representative of two independent experiments. **(E)** Relative expression of selected genes in the tongues of GF and SPF mice. Graphs present mean transcript levels ± SEM (n=6), quantified by qPCR. Data are representative of two independent experiments. **(F)** Immunofluorescence images of tongue cross-sections from GF and SPF mice stained for GAS6 (red) and DAPI (blue). Graphs display mean fluorescence intensity ± SEM (n=8). Data are representative of two independent experiments. Scale bar, 50 µm. **(G)** Flow cytometry plots and graphs showing mean frequencies ± SEM (n=4-5) of IL-1β-producing LCs in the tongue epithelium of GF and SPF mice. Data are representative of two independent experiments. **(H)** Flow cytometry histogram and graphs presenting mean frequencies ± SEM (n=3-5) of IL-1β-producing BM-derived LCs upon LPS exposure. *, *P* < 0.05; **, *P* < 0.01; ***, *P* < 0.001.

## Discussion

This study uncovers neuroimmune mechanisms that govern innervation within the tongue epithelium, emphasizing the role of LCs in promoting neuronal intraepithelial innervation. This finding supports the notion of the co-evolution of the nerve and immune systems in barrier tissues (Deng et al., 2024). The mechanism involves the production of IL-1β by the LC which induces basal epithelial cells to upregulate NGF expression, a process influenced by age and the microbiota. Whether similar mechanisms operate in the skin, where LCs also regulate epidermal innervation, remains uncertain (Zhang et al., 2021). In the skin, LCs can be categorized into two subsets based on their functions and responses: one subset is primarily active under quiescent conditions, fulfilling classical LC roles, while another predominates during inflammation and exhibits regulatory functions (Liu et al., 2021). Notably, the quiescent LC subset is the primary producer of IL-1β, though its role in local innervation has yet to be clarified. In the oral mucosa, LCs comprise distinct subsets, including pre-DC-derived LC1 and LC2 populations, as well as the third population of moLCs (Capucha et al., 2015). Our current findings, alongside previous data (Saba et al., 2022), indicate that LC2s and moLCs are the dominant subsets in the tongue epithelium, with moLCs appearing to be the main producers of IL-1β. Notably, EpCAM⁺ DCs, which serve as immediate precursors to oral LCs, represent an additional source of IL-1β in the epithelium. This may explain why the depletion of LCs does not lead to complete ablation of epithelial innervation, as EpCAM⁺ DCs are constantly present in the tongue epithelium and remain unaffected in Langerin-DTR mice. However, the precise contribution of the various LC subsets and EpCAM⁺ DCs to oral innervation requires further investigation to fully elucidate their distinct functions.

Aging is associated with a decline in the frequency of oral LCs in the gingiva, which is driven by tissue-specific factors rather than the quality of BM precursors in aged mice (Horev et al., 2020). These factors result in heightened local inflammation despite a largely unchanged microbiota in the aged host. Aging has been shown to disrupt epithelial barriers across multiple organs, contributing to dysregulated homeostasis (inflammation), lipid metabolism, and oxidative stress (Parrish, 2017). Consistently, RNAseq analysis of aged tongue epithelium revealed a compromised cellular environment, marked by reduced metabolic activity and stress-handling capacities. Notably, some of the pathways downregulated in the aged tongue epithelium are critical for LC differentiation, such as mTORC1 (Kellersch and Brocker, 2013) and OXPHOS signaling (Saba et al., 2022; Wculek et al., 2023). These findings suggest that intraepithelial innervation may be diminished in aged tissues due to the impaired ability of the tongue epithelium to support the differentiation and maintenance of LCs.

The frequencies of tongue LCs were not reduced in GF mice, contrasting with the reduction observed in the gingiva (Capucha et al., 2018; Jaber et al., 2023). Yet, the reduction in gingival LCs was primarily localized to the junctional epithelium adjacent to the dental plaque, with minimal impact on more distal gingival regions, likely explaining the absence of a similar effect in the tongue. The absence of microbiota, nevertheless, diminished IL-1β production by tongue LCs – a phenomenon also observed in mucosal CX_3_CR1^+^ mononuclear phagocytes, in which IL-1β is crucial for barrier repair (Wu et al., 2022). In the tongue, CX_3_CR1 is expressed by both LCs, the main IL-1β producer in the epithelium, and macrophages that also produce IL-1β and contribute to innervation in the lamina propria (Lyras et al., 2022). Despite this, our analysis did not detect a significant reduction in lamina propria innervation, although a strong trend was noted. This may be due to other mononuclear phagocytes present in the lamina propria secreting IL-1β, such as Fn1^+^ APCs, or the involvement of IL-1β-independent mechanisms mediating local innervation. Regarding Fn1^+^ cells, in addition to modulating IL-1β expression in human mononuclear cells (Graves and Roman, 1996), fibronectin is a potent promoter of peripheral neurite outgrowth (Lefcort et al., 1992). This dual role suggests that fibronectin might contribute to maintaining innervation in the lamina propria under GF conditions.

Besides affecting leukocyte function, the microbiota also regulates the maturation of the oral epithelium after birth (Koren et al., 2021). This includes the upregulation of various immunological and functional pathways, such as TLR2 and TLR4 singling. These might affect the capability of epithelial cells to secrete NGF, as TLR2, which is downregulated in the tongues of GF mice, was reported to regulate NGF expression in another setting (Krock et al., 2016). It is also established that the gut microbiota plays an important role in peripheral nerve development and vice versa (Cescon et al., 2024; Griffiths et al., 2024). These reports suggest that besides regulating LCs directly, the oral microbiota might also regulate neuronal-related activity by affecting tongue epithelial cells.

While the reduced sensitivity to capsaicin in LC-depleted mice demonstrates a link between oral LCs and pain perception, oral LCs may also play a role in sensing mechanical stress. A recent study highlights the critical role of LCs in detecting epithelial damage induced by early-life masticatory forces (Jaber et al., 2023). In the skin, LC depletion has decreased cutaneous innervation density and mechanical sensitivity in the footpad (Doss and Smith, 2014), suggesting that the intraepithelial innervation promoted by oral LCs may also contribute to mechanical sensing. These observations underscore the broader role of LCs in regulating sensory responses to diverse stimuli.

In summary, this study reveals a key neuroimmune role of oral LCs in promoting intraepithelial innervation in the tongue through IL-1β-driven NGF production by basal epithelial cells. Future studies should investigate the precise contributions of LC subsets and assess whether similar processes occur in humans, potentially opening new research and therapeutic avenues for oral sensory and mucosal pathologies.

**Figure S1:** Epithelial localization of tongue LCs

Flow cytometry plots and graphs depicting the mean frequencies ± SEM (n=3) of LCs in the epithelium and lamina propria of the tongue. The data shown are representative of two independent experiments.

**Figure S2:** BrdU incorporation kinetics of tongue and gingival LCs

BrdU-labeling kinetics of LCs during continuous exposure to BrdU. Representative data of one out of two independent experiments are shown as mean ± SEM (n = 3).

**Video 1:** Representative behavior of PBS-treated mice consuming capsaicin-supplemented water

**Video 2:** Representative behavior of DT-treated mice consuming capsaicin-supplemented water

## STAR Methods

## KEY RESOURCES TABLE

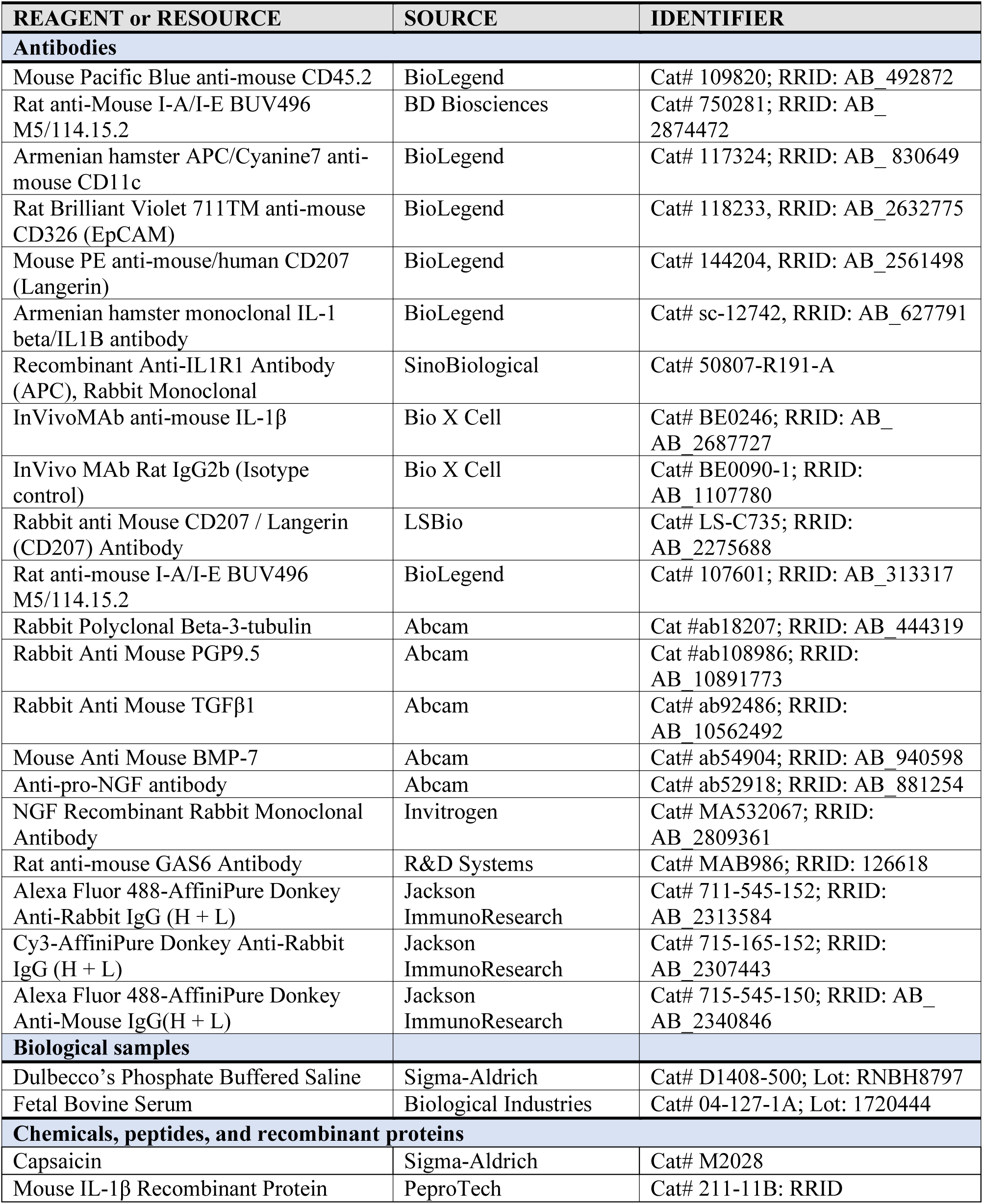
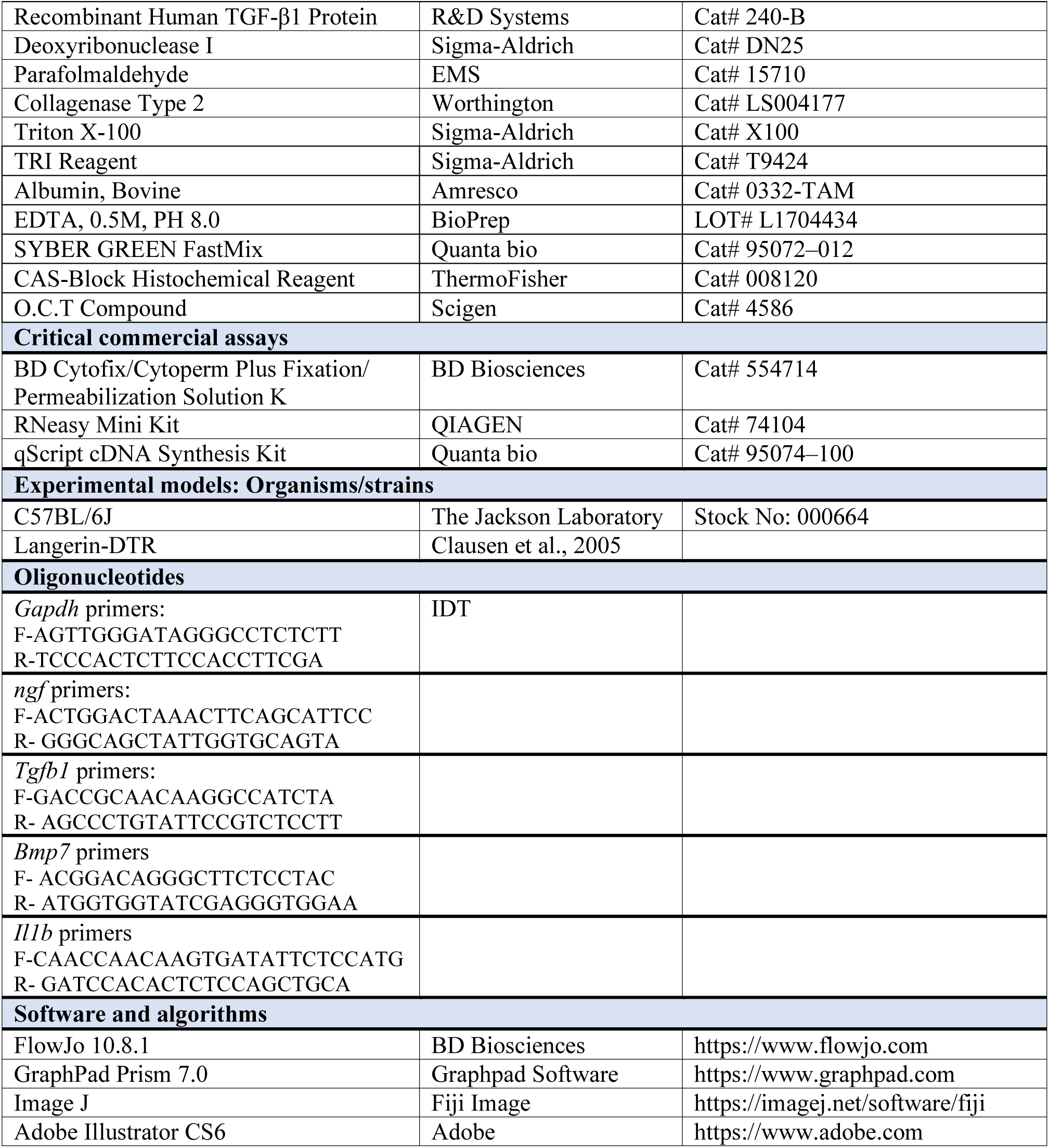

## RESOURCE AVAILABILITY

### Lead Contact

Further information and requests for resources and reagents should be directed to and will be fulfilled by the Lead Contact, Avi-Hai Hovav

### Materials Availability

This study did not generate new unique reagents.

### Data and Code Availability

The RNAseq data generated during this study will be available at GEO/NCBI after publication.

## Acknowledgments

The authors disclosed receipt of the following financial support for the research, authorship, and/or publication of this article: This work was supported by Israel Science Foundation grant 2272/20 (A.-H.H) and the German Science Foundation (DFG, Deutsche Forschungsgemeinschaft) grant CL419/2-2 (B.E.C.).

## Experimental Model and Subject Details

### Mice

C57BL/6 (B6) mice and Langerin-DTR were bred and maintained in the central animal facility at the Hebrew University Faculty of Medicine (Jerusalem, Israel). The mice were maintained under SPF conditions and analyzed at various ages as described in the text. All animal protocols were approved by the Hebrew University Institutional Animal Care and Use Committee (IACUC). The Germ-free (GF) B6 mice were maintained in sterile isolators at the Weizmann Institute of Science, the GF studies were approved by the IACUC of the Weizmann Institute of Science.

### Tongue processing

Mice were sacrificed and the tongue was excised and injected at several sites along the tongue with 0.5 mL of 4mg/ml Dispase in PBS + 2% FCS until fully distended. After incubation in PBS + 2% FCS for 10 min at 37°C the epithelium was carefully separated using forceps and then rinsed with PBS. Epithelial tissues were then minced and treated with a Collagenase type II (2 mg/mL; Worthington Biochemicals) and DNase I (1 mg/mL; Sigma) solution in PBS plus 2% FCS for 25 min at 37 °C in a shaker bath. A total of 20 μL of 0.5 M EDTA per 2 mL sample was added to the digested tissues and incubated for an additional 10 min. The cells were washed, filtered with 70-μM filter, stained with antibodies, run in Aurora (Cytek) flow cytometer, and further analyzed using FlowJo software (Tree Star).

### Immunofluorescence staining

For frozen section staining, the tongue was fixed for 2 hours in 4% PFA at 4°C, cryoprotected with 30% sucrose, and then embedded with OCT in plastic cryomolds. 30μm sections were cut from the blocks in a cryostat, collected, and immuno-stained as floating sections. The cross-sections were washed 3 times in PBS, blocked in blocking buffer (15% FBS, 5% BSA and 0.3% Triton X-100 in PBS) for 3 h at room temperature, and incubated with a primary antibody overnight at 4°C. Following 3 washing steps in PBS, the samples were incubated with a secondary antibody diluted 1:200 in blocking buffer for 2 h at room temperature, washed 3 times, stained with DAPI, and mounted. For paraffin sections, the tongue was fixed overnight at 4°C in 4% at 4°C, the tissues were subjected to gradient dehydration and paraffin embedding. Next, the tissues were micro-sectioned into 10μm-thick sections. Slides were deparaffinized with xylene, and 100%, 95%, 80%, and 70% ethanol washed 3 times with PBS. For pro-NGF and NGF staining, antigen retrieval was performed in a pressure cooker in a sodium citrate buffer at 125°C. Slides were blocked in blocking buffer (PBS, 10%FCS, 10% BSA, Triton X-100) for 1 h at room temperature and incubated with primary antibodies at 4°C. Following 3 washing steps in PBS, the samples were incubated with secondary antibodies diluted 1:200 in blocking buffer for 1.5h at RT, washed 3 times, stained with DAPI, and mounted. As a negative staining control, the primary antibody was omitted and replaced by blocking buffer. Signals were visualized and digital images were obtained using a Nikon spinning disk confocal microscope.

### Analysis of tongue intraepithelial innervation

Quantification of intraepithelial innervation was performed following the protocol by Dey *et al*. (Dey et al., 2023). Briefly, images were processed in ImageJ and converted to binary using bimodal thresholding, where the axon terminals were assigned a value of one and the background a value of zero, based on the image histogram. The number of pixels corresponding to axons was calculated. To normalize the axon value, the area of the epithelium spanned by terminal branches was measured. Innervation index values were obtained from the analyzed sections, and the average value for each animal was calculated. These averaged values were used as individual data points in the graphs, with a MATLAB script developed for batch processing.

### Ablation of langerin-expressing cells *in vivo*

Langerin-DTR mice received an intraperitoneal injection of 1µg diphtheria toxin (DT; Sigma-Aldrich) in 150 µl PBS per mouse. Additional injections of the same dose were given twice a week for the duration specified in the text.

### BrdU Incorporation Assays

8 week-old mice were injected intraperitoneally with BrdU (2 mg/mouse; Sigma) and subsequently received BrdU (0.8 mg/ml) in autoclaved drinking water changed every other day. Tissues were prepared for flow cytometric analysis as described in the previous section, and intracellular staining for BrdU was performed with the BrdU Flow kit (BD) according to the manufacturer’s protocol.

### Antibody-mediated depletion of IL-1**β**

To deplete IL-1β *in vivo*, B6 mice were administered intraperitoneal injections of 100 μg monoclonal anti-mouse IL-1β antibody (clone B122; BioXcell) in saline, twice a week from 2 to 8 weeks of age. Control mice received similar injections of rat IgG2a isotype control (anti-Trinitrophenol, clone 2A3; BioXcell).

### LC-like cell differentiation cultures

Total BM cells were isolated as previously described (Nassar et al., 2017). BM cells (5 × 10^5^ cells) were cultured in 24-well plates (Falcon) in RPMI media (450 ml RPMI 1640, 5 ml l-glutamine, 50 µM β-mercaptoethanol, 100 U/ml penicillin,100 µg/ml streptomycin, and 50 µg/ml gentamicin) supplemented with 100 ng/ml GM-CSF with or without 10 ng/ml TGF-β1 for 5 d to induce their differentiation into LC-like cells. In some experiments, 1 µg/ml LPS was added on the fifth day to stimulate the cells.

### Primary culture of tongue epithelial tissue

Mice were sacrificed and the tongue epithelium was excised as depicted above. Epithelial tissues were then equally segmented into two pieces, minced, and cultured in 24-well plates in RPMI media (450 ml RPMI 1640, 5 ml l-glutamine, 50 µM β-mercaptoethanol, 100 U/ml penicillin,100 µg/ml streptomycin, and 50 µg/ml gentamicin), The media was further supplemented with recombinant IL-1β (50 ng/ml) or TGF-β (10 ng/ml) for 72 h in 37 degrees and 5% CO2. To examine gene expression, total mRNA was extracted from the culture and analyzed by quantitative PCR as described in the RNA extraction section.

### Generation of chimeric mice

Adult CD45.1^+^ B6 mice were lethally irradiated with 950 rad, and 24 h later, the mice were injected intravenously with 2×10^6^ BM cells obtained from congenic CD45.2^+^ B6 animals to allow identification of donor-derived cells. The chimeric mice were analyzed 3 wk after BM reconstitution.

### Capsaicin aversion test

The aversive drinking test, used to assess mice’s response to noxious chemical stimuli, followed a protocol previously described (Caterina et al., 2000). Mice were habituated in the test room for at least 1 hour before the experiment. The test was conducted by two individuals, one administering the test and the other recording the results either by video or written notes. The capsaicin solution was prepared from a 4 mM stock solution containing 5% Tween 20 in saline. For 3 consecutive days, mice were allowed to drink from a bottle containing 0.125% sucrose in water for 3 hours per day. On the fourth day, the solution was supplemented with 1 μM capsaicin. The consumption of sucrose or capsaicin solutions during the daily 3-hour sessions was measured. Pain-related behavior was assessed by filming the mice for 6 minutes after the bottle was introduced.

### RNA extraction and quantification

For RNA isolation, the excised gingiva was homogenized in 500 µl TRI reagent (Sigma) using an electric homogenizer (IKA labortechnik), and RNA was extracted according to the manufacturer’s instructions. cDNA synthesis was performed using the qScript cDNA Synthesis Kit (Quanta-BioSciences). RT-PCR reactions (20 µL volume) were performed using Power SYBR Green PCR Master Mix (Quanta-BioSciences) and specific primers to the examined gene. The following reaction conditions were used: 10 min at 95 °C, 40 cycles of 15 s at 95 °C, and 60 s at 60 °C. The samples were normalized to 18S as control mRNA, by change in cycling threshold (ΔCT) method and calculated based on 2-ΔΔCT.

### RNAseq analysis

Raw reads were processed according to the QuantSeq User Guide recommendations, reads were trimmed at their 5’ end to remove the first 12 bases, then low quality and technical bases were removed from the 3’ end using cutadapt (version 1.12) (Marcel, 2011). Finally, low-quality reads, with more than 30 percent of the bases with quality below 20, were filtered out using the FASTX package (version 0.0.14). Processed reads were aligned against the mouse genome using TopHat (v2.1.1) (Kim et al., 2013). The genome version was GRCm38, with annotations from Ensembl release 89. Htseq-count (version 0.6.0) (Anders et al., 2015) was then used for quantification of raw counts per gene per sample, excluding short or otherwise unwanted gene types, such as rRNA or miRNA. Normalization and differential expression analysis were performed with the DESeq2 package (version 1.12.4) (Love et al., 2014). Genes with a sum of counts less than 10 over all samples were filtered out before normalization. Differential expression, comparing control (PBS) and treated (DT) groups, was calculated with default parameters, except not using the independent filtering algorithm. The statistical significance threshold was taken as an adjusted p-value (padj) less than 0.1. Exact commands with the full parameters used can be found under the GEO accession.

### Gene set enrichment analysis (GSEA)

Whole differential expression data were subjected to gene set enrichment analysis using GSEA (Subramanian et al., 2005). GSEA uses all differential expression data (cut-off independent) to determine whether *apriori–defined* sets of genes show statistically significant, concordant differences between two biological states. GSEA was run against the hallmark gene set collection from the molecular signatures database (MSigDB, v6.2, July 2018).

## Single-cell RNA sequencing analysis

Adult_sample count matrix files were downloaded from (Lyras et al., 2022) (https://www.ncbi.nlm.nih.gov/geo/query/acc.cgi?acc=GSE205162) and data has been re-analyze. DropletUtils was used to remove empty droplets with default settings. Cells were further filtered to keep cells with the number of UMIs above 500, number of genes above 250, and % of reads mapped to mitochondrial genes below 10%. Overall, 7,247 cells were analyzed. The Seurat package in R (Stuart et al., 2019) was used for downstream analysis and visualization. The expression data was normalized using Seurat’s NormalizeData function, which normalizes the feature expression measurements for each cell by the total expression, multiplies this by a scale factor (10,000), and then log-transforms the results. The top 2,000 highly variable genes were identified using Seurat’s FindVariableFeatures function with the ‘vst’ method. Potential sources of unspecific variation in the data were removed by regressing out the mitochondrial gene proportion and UMI count using linear models and finally by scaling and centering the residuals as implemented in the function “ScaleData” of the Seurat package. Principal component analysis (PCA) was performed. We selected 18 principal components (PC) for downstream analyses. Cell clusters were generated using Seurat’s unsupervised graph-based clustering functions “FindNeighbors” and “FindClusters” (resolution=0.5). UMAP was generated using the RunUMAP on the projected principal component (PC) space. Seurat’s functions FeaturePlot and DimPlot were used for visualization. Marker genes for each cluster were identified by performing differential expression between a distinct cell cluster and the cells of the other clusters with the non-parametric Wilcoxon rank sum test (Seurat’s FindAllMarkers function). Data was subset for APC cells (using gene expression and manual annotations) and the same analysis as above was applied to the subset population. Cell types were assigned manually based on the expression of classic marker genes. DE analysis between specific cell types was done using the FindMarkers function with default parameters. Go analysis on the DE genes was done using the ClusterProfiler package in R.

### Quantification and Statistical Analysis

Data are expressed as means ± SEM. Statistical tests were performed using unpaired t-test comparing two groups, and one-way ANOVA comparing more than two groups. A p-value of <0.05 was considered significant. Detailed information on the number (n) of biological samples and animals used can be found in figure legends. * P < 0.05 **, P < 0.01, and *** P < 0.00.1.

