## Supplementary figures and images for "Langerhans Cells Regulate Tongue Intraepithelial Innervation in a Microbiota- and Age-Dependent Manner"

### Fig. S1

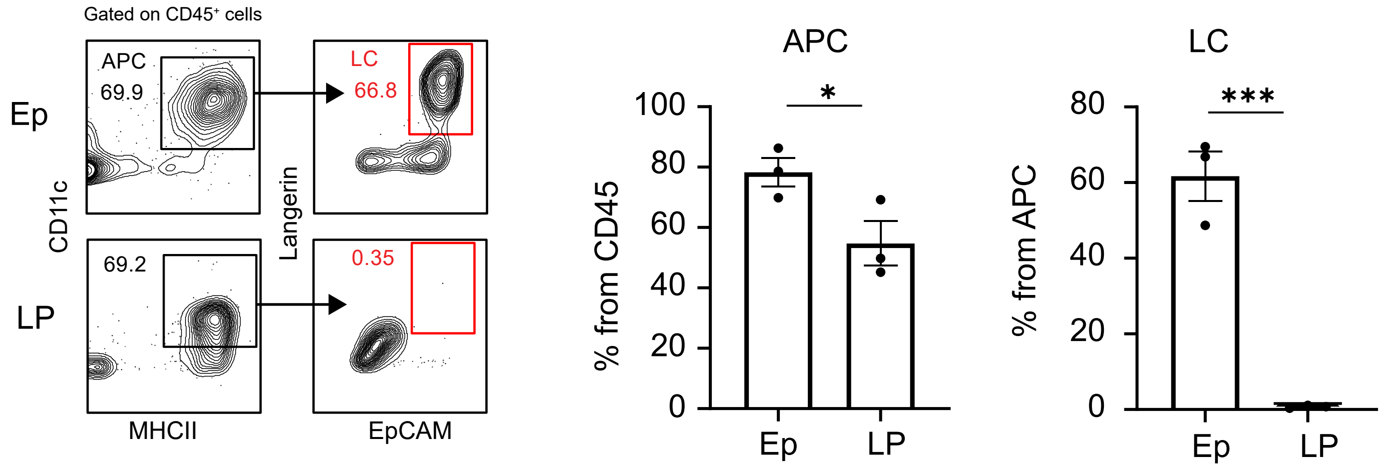

### Fig. S2

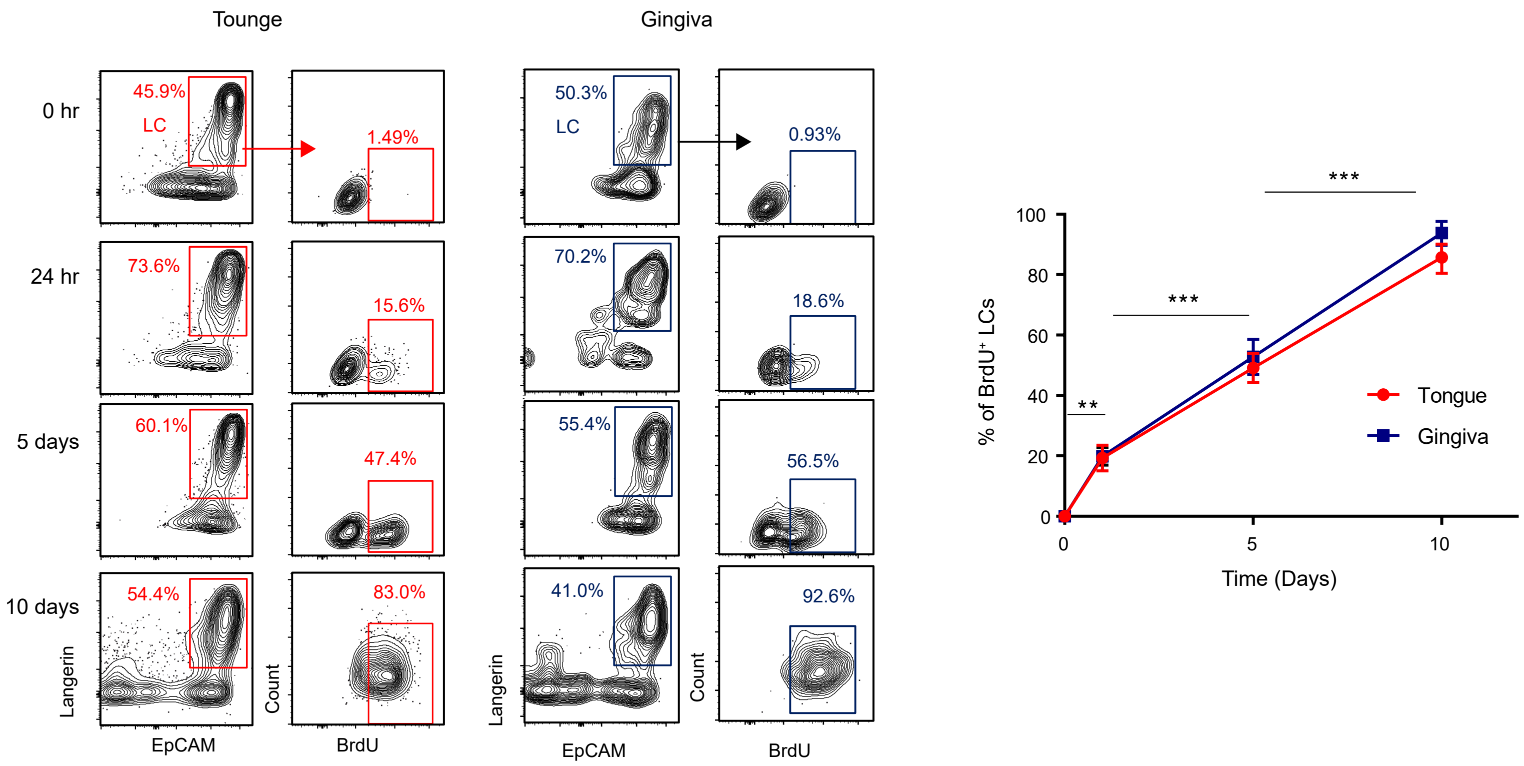
